# Comprehensive transcriptome data of normal and *Ascosphaera apis*-infected western honeybee larval guts

**DOI:** 10.1101/2020.04.11.037598

**Authors:** Yu Du, Jie Wang, Zhiwei Zhu, Haibin Jiang, Xiaoxue Fan, Yuanchan Fan, Yanzhen Zheng, Cuiling Xiong, Dafu Chen, Rui Guo

## Abstract

*Apis mellifera ligustica* is a well-known subspecies of western honeybee, *Apis mellifera. Ascosphaera apis* is a common fungal pathogen of honeybee larvae, resulting in Chalkbrood disease. In this article, deep sequencing of un-treated 4-, 5-, and 6-day-old larval guts (AmCK1, AmCK2, AmCK3) and *A. apis*-treated 4-, 5- and 6-day-old larval guts (AmT1, AmT2, AmT3) of *Apis mellifera ligustica* were conducted using Illumina HiSeq™ 4000 platform. In total, 85811046, 81962296, 85636572, 79267686, 82889882, and 100211796 raw reads were respectively yielded from above-mentioned six groups. The result of sequencing satuation analysis suggested that the sequencing depth in this work was enough to detect nearly all expressed genes. After quality control, 85739414, 81896402, 85573798, 79202304, 82828926, and 100128692 clean reads were obtained. Additionally, the GC content of each group was above 45.26%. Furthermore, 47035852, 65612676, 71803878, 62560904, 65018360, and 56278272 clean reads were mapped to the *Apis mellifera* genome, including 41221479, 61172916, 66724233, 57531335, 60245732, and 52638986 unique mapped clean reads, and 2078427, 986825, 915375, 1082925, 1097130, and 716436 multiple mapped clean reads. In addition, exons were the most abundant regions mapped by clean reads, follow by intergenic regions and introns. The strand-specific cDNA library-based RNA sequencing data documented here will faciliate study on molecualr mechanisms underlying host immune response and host-pathogen interaction during chalkbrood disease, and benefit understanding of non-coding RNA-mediated cross-kingdom regulation between *A. m. ligustica* larvae and *A. apis*.

**Value of the Data:**

- This data reported here could be used to explore circRNAs, lncRNAs, mRNAs and their competing endogenous RNA networks involved in response of western honeybee larvae to *Ascosphaera apis* infection.
- The current data contributes to better understanding mechanisms regulating host-pathogen interaction during chalkbrood disease.
- Our data can provide novel insights into understanding non-coding RNA-mediated cross-kingdom regulation between *Apis mellifera ligustica* and *A. apis*.

## 1. Data description

The share data were generated from strand-specific cDNA library-based RNA-seq of normal larval guts (AmCK1, AmCK2, and AmCK3) and *Ascosphaera apis*-infected (AmT1, AmT2, and AmT3) larval guts of *Apis mellifera ligustica*. Using next-generation sequencing, 85811046, 81962296, 85636572, 79267686, 82889882, and 100211796 raw reads were respectively gained from AmCK1, AmCK2, AmCK3, AmT1, AmT2, and AmT3 (**Table 1**). As presented in **Figure 1**, the sequencing depth was saturated and enough to detect nearly all expressed genes. Additionally, 85739414, 81896402, 85573798, 79202304, 82828926, and 100128692 clean reads were obtained from each group after quality control, and the total numbers of bases were about 12.8 Gb, 12.3 Gb, 12.8 Gb, 11.9 Gb, 12.4 Gb, and 15.0 Gb, respectively. The GC contents of clean reads from above-mentioned six groups were 49.32%, 45.26%, 46.67%, 46.19%, 46.32%, and 48.82%, respectively **(Table 2)**. Furthermore, 47035852, 65612676, 71803878, 62560904, 65018360, and 56278272 clean reads were mapped to the reference genome of *Apis mellifera*, including 41221479, 61172916, 66724233, 57531335, 60245732, and 52638986 unique mapped clean reads, and 2078427, 986825, 915375, 1082925, 1097130, and 716436 multiple mapped clean reads. As **Table 4** shown, among the mapped clean reads, 63.10%, 69.27%, 68.20%, 66.21%, 68.30%, and 69.80% were mapped to exons; 27.32%, 22.18%, 22.88%, 24.60%, 23.11%, and 20.98% to intergenic regions; and 9.59%, 8.56%, 8.92%, 9.19%, 8.59%, and 9.23% to introns.

**Table 1.** Overview of filtering of raw reads from strand-specific library-based RNA sequencing.

| Sample | Raw reads | clean reads | Adaptor | Low quality | poly A | N |
| --- | --- | --- | --- | --- | --- | --- |
| AmCK1 | 85811046 | 85739414 (99.92%) | 4080 | 65752 | 0 | 1800 |
| AmCK2 | 81962296 | 81896402 (99.92%) | 4062 | 60048 | 0 | 1784 |
| AmCK3 | 85636572 | 85573798 (99.93%) | 3530 | 57390 | 0 | 1854 |
| AmT1 | 79267686 | 79202304 (99.92%) | 4706 | 58928 | 0 | 1748 |
| AmT2 | 82889882 | 82828926 (99.93%) | 3492 | 55624 | 0 | 1840 |
| AmT3 | 100211796 | 100128692 (99.92%) | 5470 | 75566 | 0 | 2068 |

**Table 2.** Summary of clean reads after strict quality control.

| Sample | Total bases of raw reads | Total bases of clean reads | Q20 | Q30 | GC content |
| --- | --- | --- | --- | --- | --- |
| AmCK1 | 12871656900 | 12827986590 | 98.10% | 94.42% | 49.32% |
| AmCK2 | 12294344400 | 12250319245 | 98.11% | 94.36% | 45.26% |
| AmCK3 | 12845485800 | 12803375438 | 98.16% | 94.50% | 46.67% |
| AmT1 | 11890152900 | 11853296364 | 98.07% | 94.30% | 46.19% |
| AmT2 | 12433482300 | 12392905852 | 98.25% | 94.70% | 46.32% |
| AmT3 | 15031769400 | 14986997060 | 98.16% | 94.48% | 48.82% |

**Table 3.** Overview of mapping of clean reads to reference genome of *A. mellifera*.

| Sample | Total<br>clean reads | Unmapped<br>clean reads | Unique mapped<br>clean reads | Multiple mapped<br>clean reads | Mapping ratio |
| --- | --- | --- | --- | --- | --- |
| AmCK1 | 47035852 | 3735946 | 41221479 | 2078427 | 92.06% |
| AmCK2 | 65612676 | 3452935 | 61172916 | 986825 | 94.74% |
| AmCK3 | 71803878 | 4164270 | 66724233 | 915375 | 94.20% |
| AmT1 | 62560904 | 3946644 | 57531335 | 1082925 | 93.69% |
| AmT2 | 65018360 | 3675498 | 60245732 | 1097130 | 94.35% |
| AmT3 | 56278272 | 2922850 | 52638986 | 716436 | 94.81% |

**Table 4.**
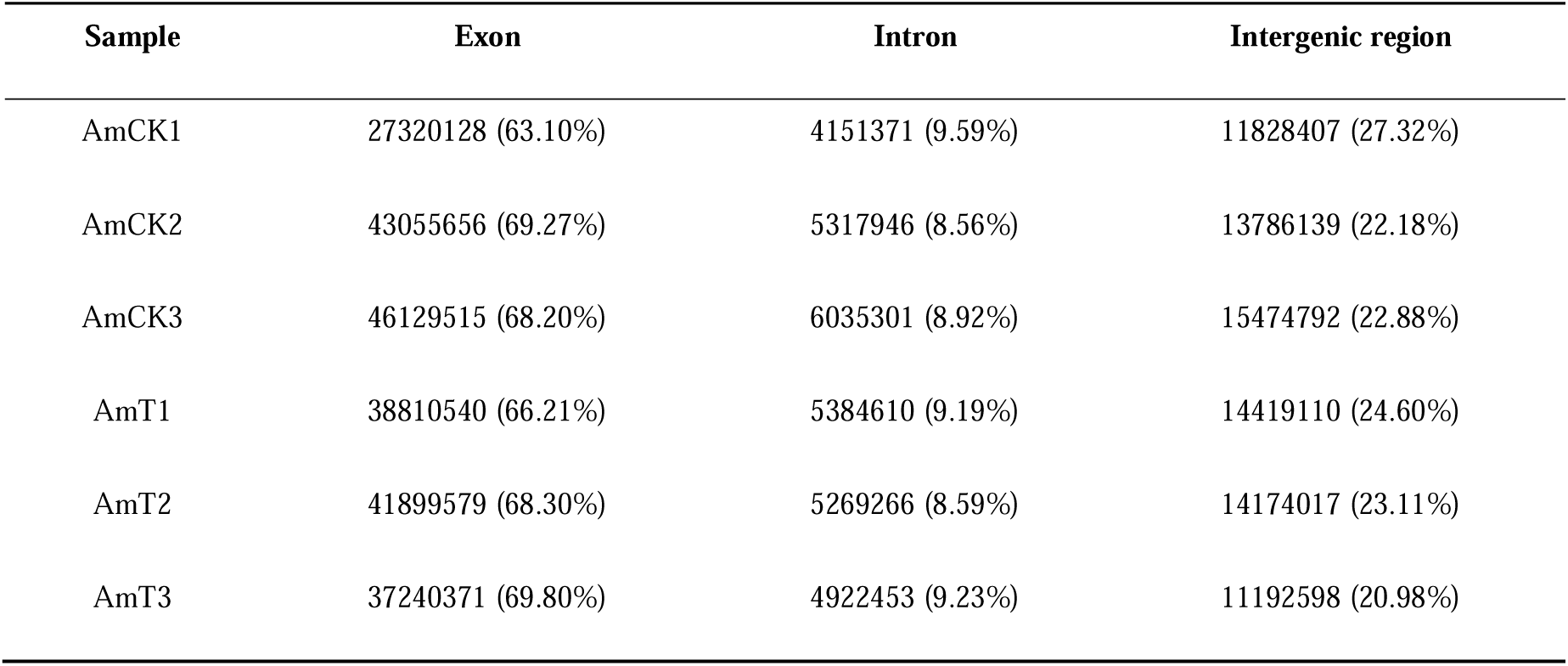
Summary of mapped region of clean reads in *A. mellifera* genome.

| Sample | Exon | Intron | Intergenic region |
| --- | --- | --- | --- |
| AmCK1 | 27320128 (63.10%) | 4151371 (9.59%) | 11828407 (27.32%) |
| AmCK2 | 43055656 (69.27%) | 5317946 (8.56%) | 13786139 (22.18%) |
| AmCK3 | 46129515 (68.20%) | 6035301 (8.92%) | 15474792 (22.88%) |
| AmT1 | 38810540 (66.21%) | 5384610 (9.19%) | 14419110 (24.60%) |
| AmT2 | 41899579 (68.30%) | 5269266 (8.59%) | 14174017 (23.11%) |
| AmT3 | 37240371 (69.80%) | 4922453 (9.23%) | 11192598 (20.98%) |

**Figure 1.**
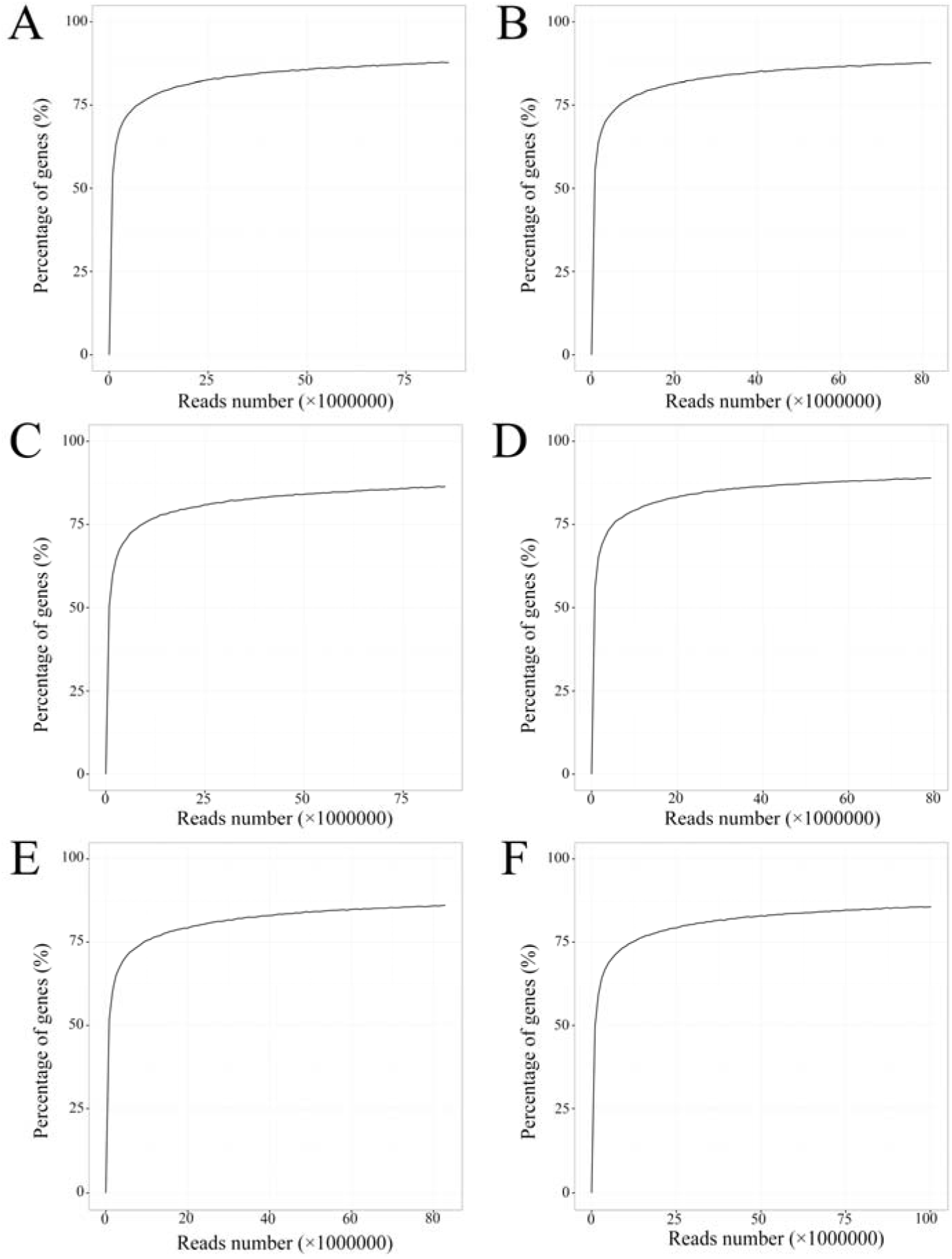
Sequencing satuation of six sample groups. A: AmCK1 group; B: AmCK2 group; C: AmCK3 group; D: AmT1 group; E: AmT2 group; F: AmT3 group

## Experimental Design, Materials, and Methods

### 2.1 Honeybee larval gut sample preparation

Following the method described by Peng et al. [1], 2-day-old larvae of *A. m. ligustica* workers were removed from the combs with a Chinese grafting tool, and carefully transferred to droplet of 10 μL artificial diet in each well of sterile 48-well culture plates. Culture plates with larvae were then incubated at 90% RH and 35±0.5 °C [2].

*A. apis* [3] was previously isolated from a fresh chalkbrood mummy following the described method by Jensen et al. [4], and conserved in Honeybee Protection Lab in College of Animal Sciences (College of bee Science), Fujian Agriculture and Forestry University. *A. apis* was cultured at 33±0.5 °C on plates of Potato dextrose agar (PDA) medium followed by purification of fungal spores at 7 d after culturing using the previously developed method [4] with some minor modifications [5]. To cause effective infection, 3-day-old larvae in *A. apis*-treated groups were fed with artificial diet containing *A. apis* spores (1×10^7^ spores/mL); while 3-day-old larvae in un-treated groups were fed with diet without spores. Following our previously developed method [2], 4-, 5- and 6-day-old larval guts from *A. apis*-treated groups and un-treated groups were respectively harvested, immediately frozen in liquid nitrogen and then kept at -80 °C until deep sequencing. *A. apis*-treated 4-, 5- and 6-day-old larval guts (n=7) were termed as AmT1, AmT2, and AmT3, while un-treated 4-, 5- and 6-day-old larval guts (n=7) were termed as AmCK1, AmCK2, AmCK3, respectively.

### 2.2. RNA extraction, strand-specific cDNA library construction and deep sequencing

Firstly, total RNA was respectively extracted from above-mentioned six groups using Trizol (Invitrogen, USA) according to the manufacturer’s instruction. RNA quality was assessed on an Agilent 2100 Bioanalyzer (Agilent Technologies, USA) and checked using 1% agarose gel eletrophoresis, respectively. Secondly, mRNAs and ncRNAs were retained after removing rRNAs and then fragmented into short fragments with fragmentation buffer (Illumina, USA) followed by reverse transcription into cDNAs with random primers. Thirdly, second-strand cDNAs were synthesized by dNTP (dUTP instead of dTTP), DNA polymerase I, RNase H, and buffer. Fourthly, the cDNA fragments were purified, end repaired, poly(A) added, and ligated to Illumina sequencing adapters with QiaQuick PCR extrAmTion kit (QIAGEN, Germany), followed by digestion of the second-strand cDNAs with UNG (Uracil-N-Glycosylase) (Illumina, USA). Finally, the digested products were size selected via agarose gel electrophoresis, PCR amplified, and sequenced using Illumina HiSeq™ 4000 platform (Illumina, USA) by Gene Denovo Biotechnology Co. (Guangzhou, China).

### 2.3. Data processing

Firstly, using fastp software [6], the generated raw reads were filtered by removing reads that contain adapters, more than 10% of unknown nucleotides (N), and more than 50% of low quality bases to gain high quality clean reads to obtain high quality clean reads. Secondly, clean reads were mapped to ribosome RNA (rRNA) database (http://rdp.cme.msu.edu/) with short reads alignment tool Bowtie2 [7]. The mapped clean reads were removed and the remaining clean reads were used for transcript assembly and following analysis. Thirdly, the rRNA-removed clean reads of each group were mapped to the reference genome of *A. mellifera* (assembly Amel_HAv3.1) using HISAT2 (version 2.1.0) following alignment parameters: (1) maximum read mismatch is two; (2) the distance between mate-pair reads is 50 bp; (3) the error of distance between mate-pair reads is ± 80 bp.

## Acknowledgments

This research was financially supported by the Earmarked Fund for China Agriculture Research System (No. CARS-44-KXJ7), the Science and Technology Planning Project of Fujian Province (No. 2018J05042), the Teaching and Scientific Research Fund of Education Department of Fujian Province (No. JAT170158), the Outstanding Scientific Research Manpower Fund of Fujian Agriculture and Forestry University (No. xjq201814), and the Scientific and Technical Innovation Fund of Fujian Agriculture and Forestry University (No. CXZX2017342, No. CXZX2017343).

